# Sex-dependent effects of peptidylarginine deiminases on neutrophil function and long-term outcomes after spinal cord injury

**DOI:** 10.1101/2025.05.08.652924

**Authors:** Shelby K. Reid, Ashley V. Tran, Miranda E. Leal-Garcia, Sachit Devaraj, Mustafa Ozturgut, Dylan A. McCreedy

## Abstract

Traumatic spinal cord injury (SCI) initiates an influx of peripheral immune cells to the spinal cord parenchyma that compound tissue damage and restrict functional recovery. Neutrophils infiltrate the spinal cord within the first day after injury, releasing extracellular traps (NETs) comprised of decondensed DNA, modified histones, and granule enzymes, that can worsen tissue damage. Peptidylarginine demininases (PADs), particularly PAD4, have been indicated as mediators of NET formation by facilitating the decondensation of nuclear chromatin via histone citrullination. Though PADs have been shown to be regulated by sex hormones, sex-differences in PAD regulation of neutrophil function in the context of CNS injury have yet to be explored. In this work, we investigated the role of PADs in recovery after SCI using Cl-amidine, a pan-PAD inhibitor. Strikingly, Cl-amidine treated mice exhibited sex-dependent changes to motor function, body weight, and white matter sparing after SCI. Acutely, Cl-amidine treated mice had reduced NET accumulation in the blood and decreased spinal cord neutrophil granularity. Analysis of publicly available scRNA-seq data revealed that female bone marrow neutrophils exhibited elevated *Padi4* expression relative to their male counterparts. We then utilized *Padi4* knockout (*Padi4*^*-/-*^) mice to assess the role of PAD4 in long-term recovery of male and female mice after SCI. While we observed no changes in motor recovery, a sex-dependent effect on tissue sparing was observed with *Padi4* deficiency. These data are the first description of sex differences in PAD-mediated neutrophil function after SCI and highlight the importance of inclusion of both sexes in pre-clinical research.

## Introduction

Traumatic spinal cord injury (SCI) is a highly debilitating condition marked by loss of motor function and sensation below the level of injury. The primary SCI event initiates a cascade of inflammatory processes that exacerbate tissue loss and impair long-term functional outcomes^1,2^. Breakdown of the blood-spinal cord barrier (BSCB) accompanies an influx of peripheral immune cells into the injured spinal cord within minutes to hours after SCI^3,4^. In the early hours after injury, neutrophils swarm to the injury site *en masse*, peaking in number in the first day after injury in rodent models of SCI^5–9^. While neutrophils have been primarily characterized as harmful in SCI, their precise role in SCI pathology and their activities in the injured spinal cord remain largely unknown.

Neutrophils are the most common circulating white blood cell population in humans and serve as patrolling first-responders to infection and injury. As critical mediators of host defense, neutrophils are equipped with a suite of effector functions including phagocytosis, release of reactive oxygen species, cytokine production, degranulation, and extracellular traps (ETs). Neutrophil extracellular traps (NETs) are a specialized effector function wherein the neutrophil will decondense its chromatin, decorate the DNA with cytotoxic granule proteins, and extrude it from the cell body into the extracellular space. ETs’ web-like structure serves to contain and kill pathogens; however, ETs can also be released in response to non-pathogens such as crystals, immune complexes, and tissue injury, causing collateral damage to host tissue^10–13^. CNS injury has been shown to stimulate ETosis, often leading to worse long-term outcomes^14–23^. In rodent models of SCI, ETosis has been linked to exacerbation of acute inflammation and worse recovery of long-term motor function^24–28^. However, sex as a biological variable has yet to be considered in ETosis after SCI; thus, any sex differences in the mechanisms of ET formation or response to treatment remain unknown.

While PAD4 citrullination of histones is thought to be the primary mediator of chromatin decondensation in NETosis, other PADs are also expressed in neutrophils and may play a redundant role to PAD4^11,29–32^. In the present study, we assessed the role of PAD activity and PAD4 in ET formation after SCI using pharmacological and genetic approaches with a particular focus on sex as a biological variable. We found that PAD activity has sex-dependent effects on long-term outcomes following SCI including motor recovery, weight loss, and white matter sparing. Acutely, we show that PADs mediate neutrophil responses in the injured spinal cord. Furthermore, using a PAD4 knockout, we show that PAD4 regulates long-term tissue damage after SCI in a sex-dependent manner. Together, these data show sex-dependent roles for PADs in acute inflammation and long-term outcomes following SCI, highlighting the necessity of including sex as a biological variable in neurotrauma research.

## Results

### PAD inhibition alters long term SCI outcomes in a sex-dependent manner

To assess whether PADs play a role in long-term outcomes following SCI, while also considering sex as a biological variable, we utilized a pharmacological approach to broadly inhibit PAD activity. Cl-amidine is a potent, irreversible small molecule pan-PAD inhibitor frequently used in animal studies to abrogate ET formation ^33,34^ (Fig 1A). Cl-amidine administration had no appreciable effect in the BMS main score (Fig 1B); however, sex-dependent effects on motor recovery were evident in the BMS subscore (Fig 1C, Time x Sex x Treatment interaction p=0.01165) with male mice trending towards improved subscores and female mice trending towards reduced subscores. Body weights of the mice were assessed throughout the course of the study to monitor the health of the animals as a sharp reduction in body weight can be indicative of poor health and require euthanasia of the animal. We observed a sex-dependent effect of Cl-amidine on body weight retention (Fig 4D, Time x Sex x Treatment, p=0.032) with male mice trending towards greater body weight retention and female mice trending towards greater weight loss. Mice exhibiting a BMS score ≥ 4 by 28 dpi were assessed via the horizontal ladder rung walking test (LRWT). However, no statistical differences were found between groups in percentage stepping or cumulative errors (Fig 1E-F).

**Figure 1:**
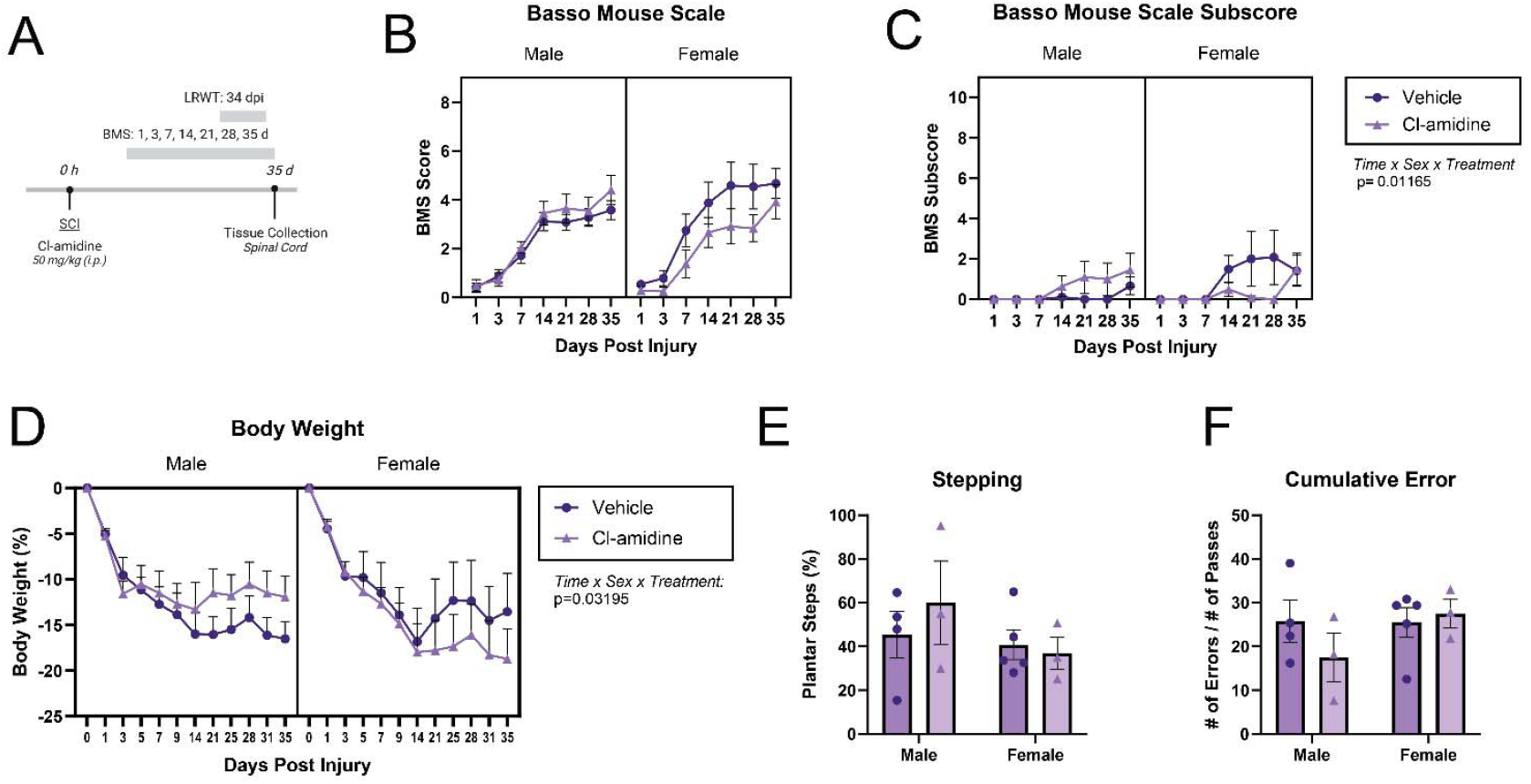
PAD inhibition with Cl-amidine alters long-term motor recovery and weight loss after SCI in a sex-dependent manner. **(A)** Cl-amidine (50 mg/kg) was administered intraperitoneally immediately following moderate T9 contusion. Functional recovery was assessed for 5 weeks following injury via the Basso Mouse Scale (BMS) and the horizonal ladder rung walking test (LRWT). **(B-C)** While no significant differences between groups were observed via the **(B)** main BMS score, **(C)** BMS subscores were differentially altered between sex and Cl-amidine treated groups across time (C: Time x Sex x Treatment, p= 0.0116; 3-way RM ANOVA, n=6-10/treatment/sex). **(D)** Body weight retention over time following SCI was differentially altered between sex and Cl-amidine treated groups. (Time x Sex x Treatment, p=0.0320; 3-way RM-ANOVA, n=6-10). **(E-F)** At 34 dpi, mice with BMS ≥ 4 were assessed via LRWT. No statistically significant differences in **(D)** percentage of plantar stepping or **(E)** cumulative error across trials were observed (2-way ANOVA, n=3-5). Data shown as mean ± SEM. *p<0.05, **p<0.01.

After tissue collection at 35 dpi, we examined white matter tissue sparing to assess the long-term tissue damage across the lesion site. We found a sex-dependent effect of Cl-amidine treatment on white matter tissue sparing (Fig 2A, Treatment x Sex, p=0.040) and lesion length (Fig 2B, Treatment x Sex, p=0.015). Specifically, we observed greater lesion length, as well as a strong trend (p=0.059) towards reduced white matter sparing, in Cl-amidine treated female mice relative to vehicle controls. No differences were observed in male mice. No effect of treatment or sex was observed for white matter sparing at the lesion center (Fig 2C). Together, these data demonstrate an interesting sex-dependent effect wherein female mice appear to be adversely impacted by PAD inhibition after SCI while male animals are either positively or not at all impacted by the same treatment.

**Figure 2:**
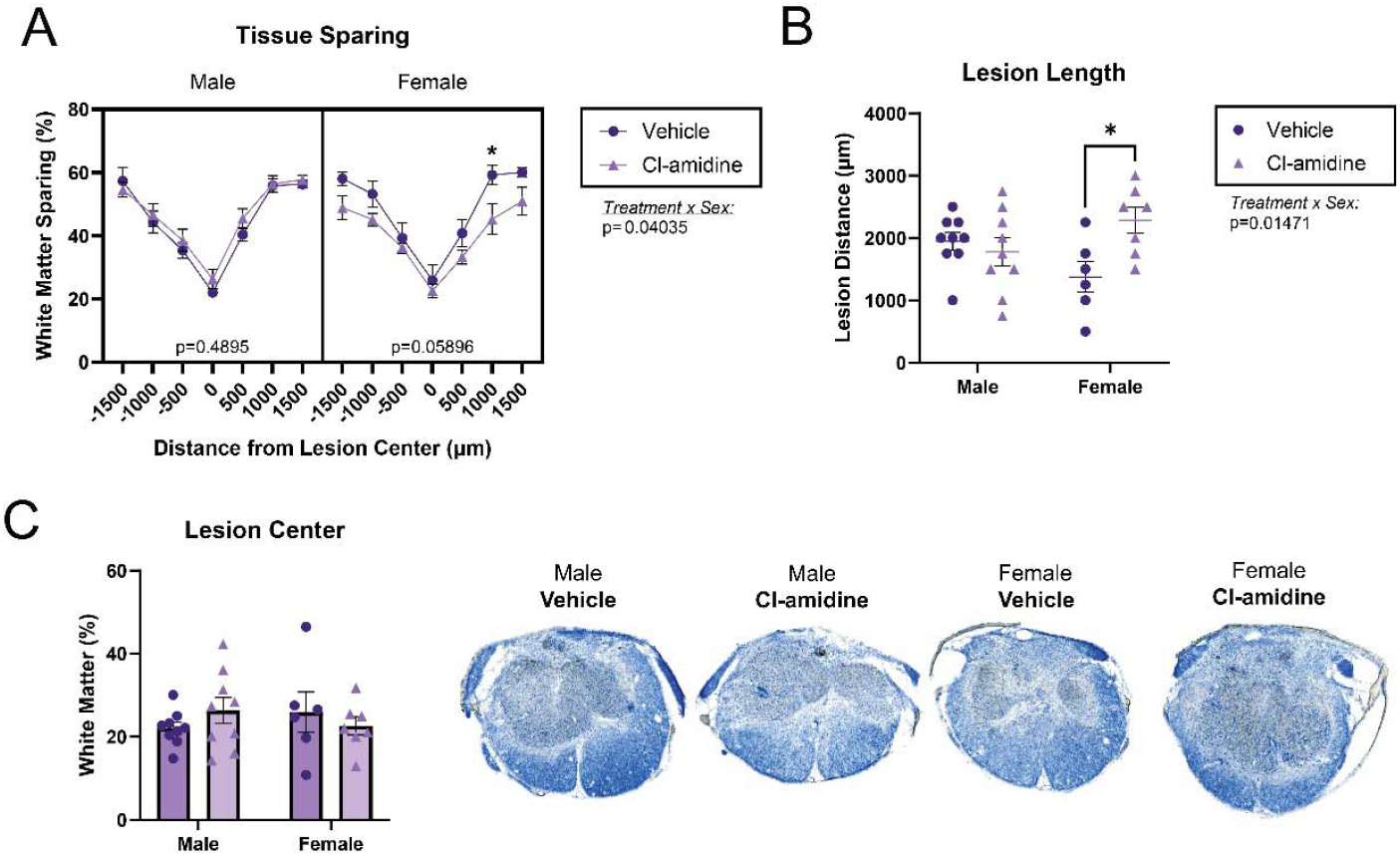
PAD activity alters long-term tissue damage after SCI in a sex-dependent manner. White matter tissue sparing in the spinal cord was assessed at 35 dpi. **(A)** Across the lesion, tissue sparing was differentially affected by sex and Cl-amidine treatment (3-way RM ANOVA, Treatment x Sex p=0.0403. Data were disaggregated by sex and performed 2-way ANOVA with Tukey’s *post-hoc*; n=6-9/sex/treatment). **(B)** Lesion length was similarly altered by treatment and sex (2-way ANOVA, Sidak’s *post-hoc*, n=6-9) with Cl-amidine increasing lesion length in female mice. **(C)** No sex or treatment effects were observed for white mater sparing at the lesion center (2-way ANOVA). Tissue images are representative of group means. Data shown as mean ± SEM. *p<0.05, **p<0.01.

### PAD inhibition reduces ET accumulation and neutrophil proportion acutely after SCI

To elucidate potential mechanisms behind the sex-dependent effects we observed in long-term recovery, we assessed Cl-amidine treated mice at 1 dpi, the peak of neutrophil infiltration and NET formation in the spinal cord^28^. We first quantified overall levels of ET complexes in cell-free supernatant of the spinal cord homogenate via capture ELISA. At 24 hpi, Cl-amidine treated mice had reduced ET-complex load in the blood (Fig 3A) with a similar trend in the spinal cord (Fig 3A).

**Figure 3:**
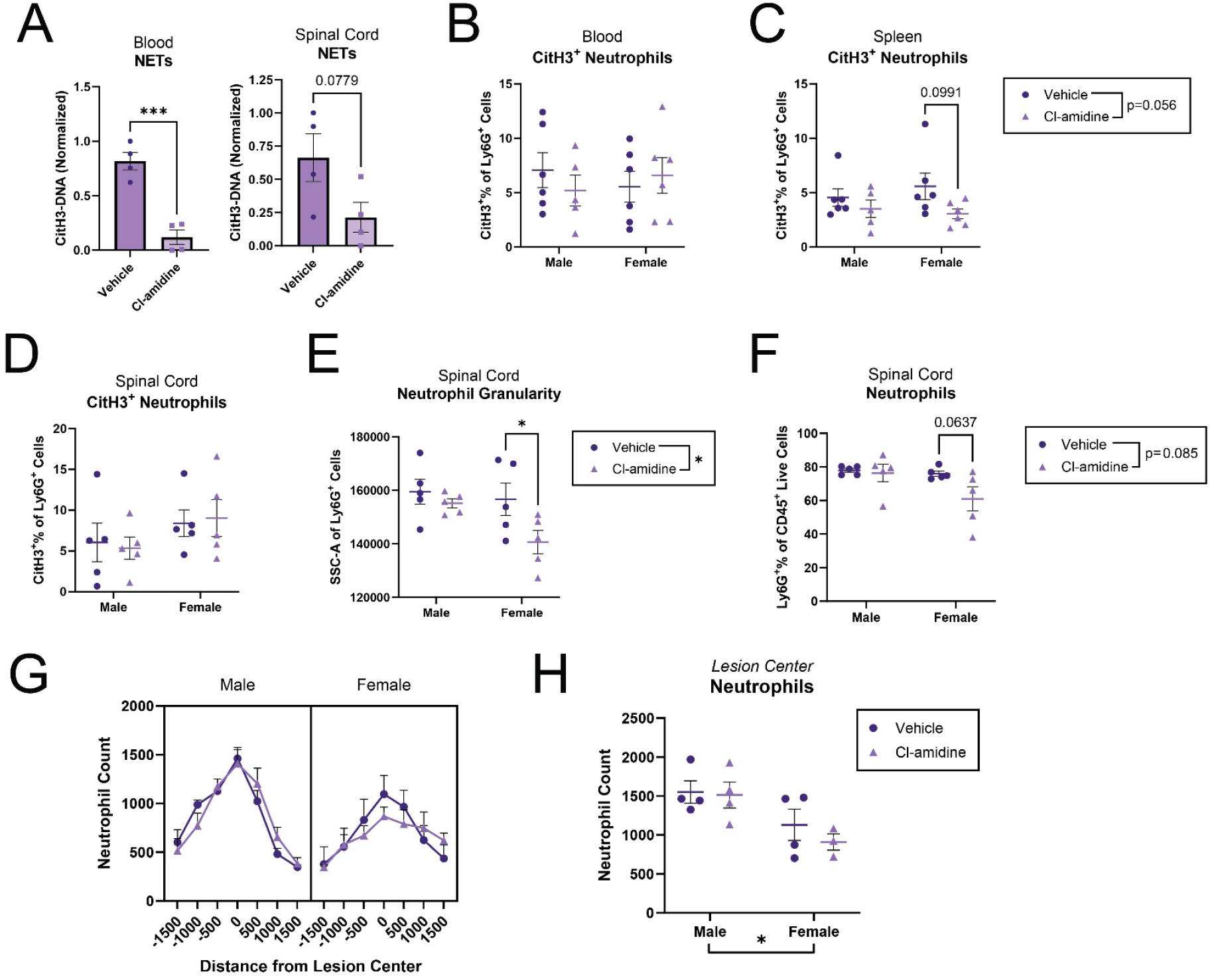
PAD inhibition with Cl-amidine reduces ET trap formation and alters immune cell response post-SCI. Immediately following moderate T9 contusion (0 hpi), Cl-amidine (50 mg/kg) was administered via intraperitoneal injection. Tissues were collected at 24 hpi. **(A)** Cl-amidine administration reduced ET complex accumulation in the blood compared to vehicle controls. A similar strong trend was observed in the spinal cord (Unpaired Student’s t-test, p_SpinalCord_=0.0779). **(B-D)** Percentage of neutrophils that were CitH3+ were assessed via flow cytometry. **(B)** Blood neutrophils showed no differences across groups while **(C)** spleen neutrophils from Cl-amidine treated animals exhibited a strong trend towards reduced CitH3 accumulation. **(D)** No differences were observed in CitH3+ neutrophils in the spinal cord (2-way ANOVA, Sidak’s *post-hoc)*. **(E)** Cl-amidine treatment differentially altered spinal neutrophil granularity by sex, (Treatment x Sex, p=0.002065. Grubbs’ test, 2-way ANOVA, Sidak’s *post-hoc*). (**F)** A strong trend towards reduction in the neutrophil proportion of CD45+ immune cells in the spinal cord was observed in female Cl-amidine treated mice (Treatment: p=0.008458; 2-way ANOVA, Sidak’s *post-hoc*). **(G-H)** No treatment group differences were observed in total number of neutrophils, though females were observed to accumulate fewer neutrophils in the spinal cord at 24 hpi (G: 3-way RM ANOVA, Tukey’s *post-hoc*. H: 2-way ANOVA). Data shown as mean ± SEM. *p<0.05, **p<0.01, ***p<0.001.

**Figure 4:**
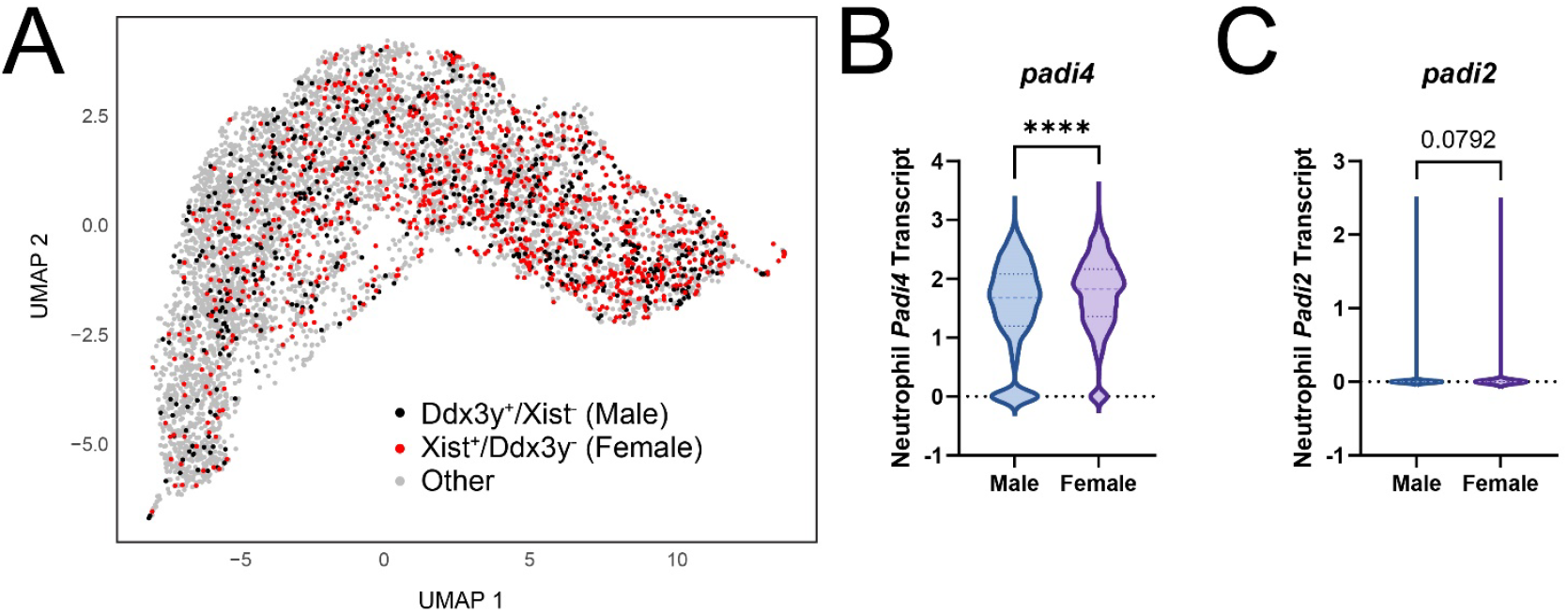
Intraspinal neutrophil PAD expression differs by sex. **(A)** UMAP of scRNAseq data for male and female bone marrow neutrophils. Male neutrophils (black) are identified by the expression of Ddx3y and female neutrophils (red) are identified by the expression of Xist. Neutrophils with no detectable expression of Ddx3y or Xist are shown in grey. **(B-C)** Comparison of *Padi4* **(B)** and *Padi2* **(C)** transcript levels between male (Ddx3y^+^/Xist^-^) and female (Xist^+^/Ddx3y^-^) neutrophils. Mann-Whitney U test. ****p<0.0001.

Although neutrophils make up the bulk of infiltrating peripheral immune cells at 24 hpi, we used flow cytometry to verify cell-type specificity of NET-inhibition. While blood neutrophils exhibited no differences across groups (Fig 3B), splenic neutrophils showed a strong trend towards a reduced percentage of CitH3^+^ neutrophils in Cl-amidine treated animals, irrespective of sex (Fig 3C). Cl-amidine did not significantly alter the percentage of CitH3^+^ intraspinal neutrophils (Fig 3D), suggesting that PAD inhibition may affect other neutrophil activities. The flow cytometry side scatter (SSC) measurement reflects the structural complexity of the immune cell and can indicate the granularity of the cells’ contents as well as cell membrane roughness accompanying cytoskeletal reorganization^35–37^. In Cl-amidine treated mice, we observed reduced neutrophil granularity (p=0.0376), which was particularly pronounced in female mice (Fig 3E, p=0.045). Cl-amidine treated females also showed a trend towards reduced neutrophil proportion out of CD45^+^ immune cells (Fig 3F, p=0.064). Interestingly, when absolute neutrophil numbers were assessed across the SCI lesion via IHC, no treatment effects were observed in either sex, though female animals accumulated notably fewer neutrophils than their male counterparts (Fig 3G-H). Together, these data show that PAD inhibition alters neutrophil activity in the injured spinal cord in a sex-dependent manner and that neutrophil response to SCI differs by sex.

### PAD gene expression in neutrophils differs by sex

To determine which PADs are expressed by neutrophils, we performed an independent analysis of publicly available single cell RNA sequencing (scRNAseq) data from male and female bone marrow neutrophils^38^. Male and female neutrophils were identified by the presence of transcripts for the sex-specific genes Ddx3y and Xist, respectively. Interestingly, we found that bone marrow neutrophils from females had higher transcript levels for *Padi4* relative to males. A similar trend was observed for *Padi2*. Transcript levels for other PAD genes were minimally detected. Our findings indicate that PAD4 is most prominent in neutrophils and that expression of *Padi*4 differs between female and male neutrophils.

### PAD4 regulates tissue sparing after SCI in a sex dependent manner

To assess the specific role of PAD4 in long-term functional outcomes after SCI, we utilized a global knockout of *Padi4* (*Padi4*^*-/-*^) and assessed motor recovery for 35 dpi (Fig 5A). Surprisingly, no differences were observed between *Padi4*^*-/-*^ mice and their wild-type (WT) littermate controls in body weight retention or BMS score by 35 dpi (Fig 5B-C). However, examination of white matter sparing at the SCI lesion epicenter (Fig 5E-F) showed a significant interaction between sex and genotype, indicating that PAD4 may differentially affect tissue sparing after SCI in a sex-dependent manner. There was also a strong trend (p=0.053) towards reduced white matter sparing at the lesion epicenter in female *Padi4*^*-/-*^ mice relative to WT controls. These data indicate that while *Padi4*^*-*^ deficiency has no apparent effect on motor recovery after SCI, PAD4 may differentially affect sparing of myelinated spinal cord tissue in male and female animals.

**Figure 5:**
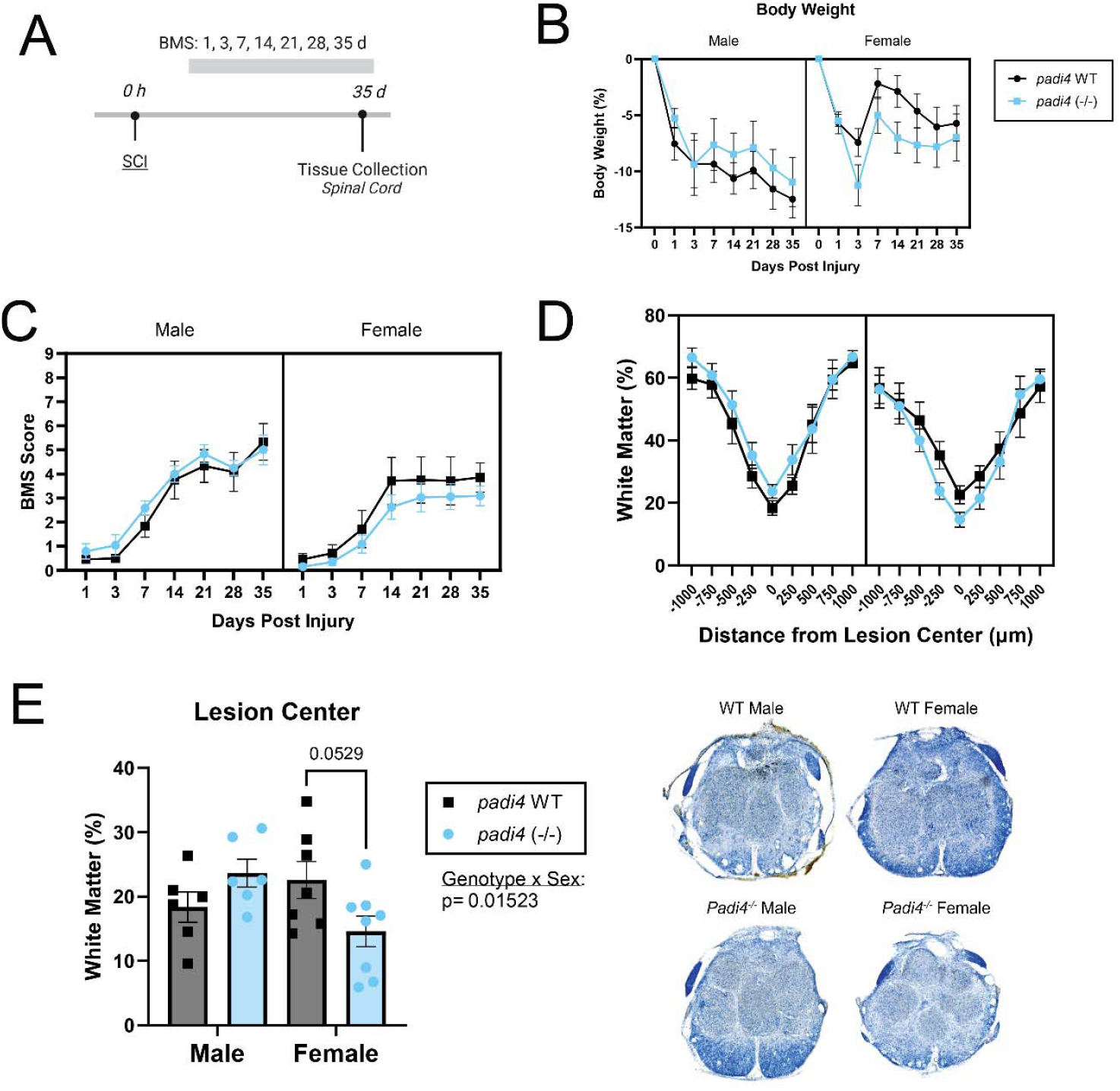
PAD4 alters tissue sparing after SCI in a sex-dependent manner. **(A)** Functional motor recovery was assessed for 5 weeks after moderate contusion SCI via the Basso Mouse Scale (BMS). **(B)** No differences in weight loss were observed between genotypes. (3-way RM ANOVA, n=6-8/sex/genotype). **(C)** No differences were observed in BMS score (3-way RM ANOVA, n=6-8/sex/genotype). **(D-E)** Spinal cord tissue centered on the lesion site was assessed for white matter tissue sparing. While no overall differences were observed **(D)** across the lesion, we observed a significant interaction between sex and genotype regarding white matter sparing at the **(E)** lesion epicenter (D: 3-way RM ANOVA (n=6-8/sex/genotype. E: 2-way ANOVA, Sidak’s *post-hoc*). Tissue images are representative of group means. Data shown as mean ± SEM.

ETs are released by neutrophils (among other cell types) in the injured spinal cord^24,39,40^, and PAD4 has been considered the canonical contributor to histone citrullination in the process of ETosis. To determine if PAD4 is involved in ETosis acutely after SCI, we assessed the abundance of ETs in the spinal cord lesion site and peripheral blood of *Padi4*^*-/-*^ mice and their wild-type littermates in a contusion model of SCI (Fig S1A). In *Padi4*^*-/-*^ mice, ET complex deposition was reduced in the spinal cord at 24 hpi (Fig S1B). Since neutrophils are most numerous in the spinal cord acutely after injury, we assessed NETs *in vivo* by flow cytometry at 24 hpi. Blood neutrophils from *Padi4*^*-/-*^ mice had significantly less intracellular histone citrullination than WT controls (Fig S1C) and intraspinal neutrophils exhibited a similar trend (Fig S1D). Interestingly, *Padi4*^*-/-*^ neutrophils from theblood had significantly higher levels of extracellular myeloperoxidase (MPO), potentially indicating degranulation, than WT controls (Fig S1E), which was not observed in the spinal cord (Fig S1F). Together, these data indicate that PAD4 mediates NET formation in SCI.

## Discussion

Extracellular traps are a double-edged sword: not only can they harm potential invaders, but also host tissues. Thus, it may not be advantageous for all cells to undergo ETosis; indeed, only up to about 25% of neutrophils will choose NETosis when faced with infection^41^. This balancing act—a cell’s propensity to choose ETosis over other potential effector functions—is dependent on many factors including pathogen size^42^, cell maturity^43,44^, age^45–48^, and sex^47,49–54^. Sex-differences in ETosis have not been widely studied; however, ETosis is thought to be directly influenced by sex hormones^49,50,54–57^. Furthermore, estrogen can directly stimulate expression of *Padi4* and *Padi2*, while androgens suppress their expression^58–63^. Despite these factors, many studies of ETosis do not include both sexes and those that do include both sexes often do not disaggregate the data by sex for subsequent statistical analyses. To the best of our knowledge, we have for the first time elucidated sex differences in the role of PAD activity on the acute inflammatory response following SCI. Interestingly, Cl-amidine administration altered the properties of neutrophils in the injured spinal cord in a sex-dependent manner. Female Cl-amidine-treated mice were observed to have reduced neutrophil proportion and neutrophil granularity compared with vehicle-treated controls. While we did not observe any treatment-dependent differences in total neutrophil accumulation across the lesion, we did observe that male animals had significantly higher neutrophil counts compared to females independent of treatment. These data indicate that composition of myeloid cell populations present after SCI may be both PAD- and sex-dependent, potentially proceeding via different mechanisms. Therefore, therapeutic interventions designed to target ETosis and ET-mediated damage must take sex-differences into account.

A common observation in both the laboratory and the clinic is that after neurotrauma, females recover better— or at least, differently—than their male counterparts^64–67^. Acute inflammation after SCI likewise differs in a sex-dependent fashion, which leads to alteration in long-term recovery outcomes^68–71^. As we observed changes to the acute inflammatory profiles of both PAD-inhibited animals, we assessed long term function by sex in both groups. Interestingly, broad PAD-inhibition with Cl-amidine resulted in sex differences body weight retention, motor recovery, and tissue sparing across the lesion. While Cl-amidine treated male mice trended towards improved long-term outcomes, Cl-amidine treated female mice were adversely affected. A potential explanation for this difference is that Cl-amidine administration inhibits PAD2, which is known to contribute to myelination and oligodendrocyte development^72^. The mechanism underlying the selective vulnerability of female animals to PAD inhibition after SCI remains a question for future study.

PAD4 has been heavily implicated in ET formation and is considered the primary enzyme responsible for histone citrullination in the canonical pathway of ET formation^73–75^. However, PAD4-independent NET formation has been reported^73,76–82^. PAD2 is also expressed in neutrophils can mainly be found in the cytosol, though it can relocate to the nucleus in response to calcium signaling^29,83,84^. While PAD2 is not required for ETosis^85,86^, recent studies have indicated that PAD2 may play a redundant role to PAD4 in ET formation^30,31,58,87–90^. Here, we confirm that *Padi4* levels differ between male and female neutrophils, at least naïve bone marrow populations. *Padi4* deletion in our model of injury did not significantly alter body weight, or motor recovery following SCI. However, white matter tissue sparing displayed a sex and genotype-dependent effect at the lesion epicenter. We also found that PAD4 is responsible for a portion of ETosis in SCI and that PAD4 mediates NETs formation in both the blood and spinal cord following injury. Interestingly, we also saw an increase in extracellular MPO on neutrophils in the blood, perhaps indicating that—without PAD4, neutrophils may choose to undergo degranulation instead of ETosis. Surprisingly, pan-PAD inhibition with Cl-amidine had no effect on NET formation (CitH3^+^ neutrophils), albeit overall ET levels were reduced. It is possible that broad inhibition of PADs primes neutrophils for PAD-independent NETosis.

In the present study, we demonstrate sex differences in PAD-mediated acute inflammation and long-term recovery in a mouse model of SCI. Immune cell accumulation in the spinal cord was PAD and sex-dependent, with PAD activity affecting neutrophil proportion in female mice only. Long-term functional and tissue sparing outcomes after injury were altered in a sex- and PAD-dependent manner. These results suggest that while PADs mediate ETosis, some PAD activity is required for optimal recovery from SCI in female animals, underlining the cruciality of assessing sex differences at the pre-clinical stage.

## Methods

### Study Design

The objective of this study was to determine whether PAD4 contributes to acute extracellular trap formation and the progression of pathology (tissue sparing, motor recovery) after spinal cord injury with a focus on unveiling any sex-dependent differences. All procedures were carried out in compliance with Texas A&M University Institutional Animal Care and Use Committee guidelines (approval numbers 2019-0045 and 2021-0340).

### Mice

C57Bl/6J mice (3-6 months) obtained from Jackson Laboratory and maintained in house were used for all pharmacological studies (#000664). For genetic studies, B6.Cg-*Padi4*^*tm1*.*1Kmow*^/J (*Pad4*^-^, #:030315) were obtained from Jackson Laboratory and maintained in house. *Pad4*^-/-^ mice lack exons 9 and 10 essential to the PAD4 active site as well as 4 exons related to Ca^2+^ binding^91^. Animals were housed in conventional 12-hr light-dark cycle with food and water accessible *ad-libitum* in cages containing 2-5 mice. Following surgery, male mice were singly housed to prevent incidents of aggression and were given huts for enrichment while female mice were housed together in groups of 2-3 per cage.

### Spinal Cord Injury

#### Surgery

Mice were anesthetized with 2% isoflurane for 10 minutes prior to surgery and kept under isoflurane for the entirety of the procedure. Following toe pinch to ensure deep anesthetization, mice were shaved and prepared with betadine and 70% ethanol to sterilize the incision area. The spinal column over the thoracic vertebral level nine (T9) lamina was exposed, the lamina was removed, and the spinal column was clamped at T8 and T10. A 60 kDyne injury with a 1 second dwell time was applied to the exposed spinal cord at T9 via an Infinite Horizons / Precision Systems and Instrumentation impactor. Superficial muscle was closed with a 5.0 suture and 50 µL of bupivacaine was administered topically to mitigate post-operative pain. The skin was closed with wound clips and given 1.0 mL of subcutaneous saline to prevent dehydration. Mice were allowed to recover in a home cage warmed on a heating pad until they recovered from anesthesia.

#### Post-Operative Care

Mouse bladders were manually voided twice per day for the entirety of the experiment. To prevent urinary tract infection and dehydration, mice were administered Cefazolin (120 mg/kg in ∼1 mL of saline) and 1.0 mL of saline daily for 10 days. Weights were assessed daily for 10 days and weekly thereafter for the entirety of the study.

### Cl-amidine Administration

To inhibit PAD activity pharmacologically, Cl-amidine (Sigma-Aldrich) was utilized. Immediately following the conclusion of SCI surgery, Cl-amidine was administered via intraperitoneal injection at 50 mg/kg while control animals received vehicle only. For Figures 3.3A and 3.3C, Cl-amidine was dissolved in 100% DMSO. For all other figures, 0.5% DMSO in saline was used as the delivery vehicle. Experimenters were blinded to treatment groups and treatment groups were randomized prior to the study.

### Motor Recovery

#### Basso Mouse Scale

Following surgery, mice were assessed at 1 and 3 dpi and weekly thereafter for 35 dpi via the Basso Mouse Scale (BMS). Assessment was completed at least 1 hour post bladder voiding and carried out in room mice were housed in. BMS assessment was completed by 2 independent raters during a 4 minute period in which the mouse was allowed to roam freely in a clear acrylic open field with a mirror placed underneath. Scores were averaged between the independent raters. Animals with BMS > 2 at 1 dpi were excluded from the study.

#### Ladder Rung Walking Test

Mouse Horizontal Ladder (Maze Engineers) was used to assess coordinated motor function in mice that had achieved BMS ≥ 4 by 28 dpi. Ladder was mounted across an acrylic bin equipped with strip LED lighting to illuminate the walkway. A ramp at the end of the ladder extended into the mouse’s home cage. Mirrors were mounted at an approximately 45° angle to the walkway such that the mouse was simultaneously visible from beneath and the sides. A GoPro Hero9 was mounted below the walkway to capture footage in 4K at 60 FPS using the Linear FOV. Mice were acclimated to the ladder and ramp in their home cage for 3 min, then placed on the side of the ladder opposite to their home cage and allowed to cross and descend a ramp into their home cage. Mice were allowed to rest for 2 minutes in home cage between trials.

This process was repeated until the mouse voluntarily exited the ladder. Mice were trained for 3 days prior to final assessment, which took place at 35 dpi. Mice were assessed for 5 passes. Videos were assessed by a blinded rater^92^. Briefly, each step was scored as plantar, toe, skip, slip, miss, or drag for both left and right hind paws. Percent stepping was quantified as Plantar + Toe steps / All steps x 100. Stepping errors were quantified as total Slips + Skips + Miss + Drag / number of passes.

### Tissue Processing

#### Histology

Tissues were collected as previously described^28^. Briefly, mice were administered a lethal dose of 2.5% Avertin then transcardially perfused with 25 mL of cold 1X PBS followed by 25 mL of cold 4% paraformaldehyde to fix tissues. Spinal columns were then dissected and left in 4% paraformaldehyde to fix overnight. Spinal cords were then dissected and cryoprotected in 30% sucrose for 2 days. Following this, 4 mm of spinal cord was mounted in OCT and cut in 25 µm sections, then stored at -80C.

#### Flow Cytometry / ELISA

Tissues were collected as previously described^28^. Briefly, mice were administered a lethal dose of 2.5% Avertin via intraperitoneal injection. Following euthanasia, blood was drawn from the heart in an EDTA coated 1 mL syringe using a 25 G needle. The needle was removed from the syringe and blood was ejected into a chilled microcentrifuge tube (MCT) then diluted with an equal volume of 2.5 mM EDTA in HBSS. MCT was inverted 8-10 times to mix and immediately centrifuged at 1350 G for 5 min at 4C. Supernatant was transferred to a fresh chilled MCT and centrifuged (1350 G, 5 min, 4C), then aliquoted and stored at -80C for ELISA. Pellet was transferred to 10 mL of ice-cold HBSS containing 50 µL of 0.5 M EDTA and kept on ice. Spinal cord (5 mm) was rapidly dissected and placed into 300 µL of ice-cold RPMI and kept on ice. Spleen was collected and connected fat was removed. Excess moisture was removed with a Kim Wipe and the spleen was weighed before being transferred to 300 µL of ice-cold RPMI.

### Flow Cytometry

Cells were treated as previously described (Reid *et al*., 2025). All steps were taken on ice and with ice-cold solutions. Briefly, cells underwent RBC lysis (BD PharmLyse solution, 1:10 dilution in water, BD Biosciences). Blood was incubated for 6 min with 10 mL of solution while spinal cord and spleen were incubated for 5 min with 1 mL of solution. Samples were subsequently diluted with NDS-FACS buffer (2% Normal Donkey Serum (Lampire), 1mM EDTA (Invitrogen) in HBSS without Ca^2+^, Mg^2+^ and Phenol Red (Corning)) - 40 mL for blood, 25 mL for spleen, 10 mL for spinal cord. Samples were centrifuged (400 RCF, 5 min, 4°C; settings used for all remaining centrifuge steps), resuspended in HBSS, then centrifuged. Total spleen cell counts were assessed using the EVE Automated Cell Counter (NanoEntek) according to manufacturer’s instructions. Pellets were resuspended in Zombie Red Fixable Viability Dye (1:500 in 100 µL of HBSS, BioLegend) according to manufacturer’s instructions and incubated for 30 min. Cells were washed with NDS-FACS, centrifuged, then transferred to a 96-well plate. Following resuspension in NDS-FACS and centrifugation, cells were blocked with anti-CD16/32 antibody (clone 93, BioLegend, 1:100) for 20 min. Cells were diluted with NDS-FACS buffer and centrifuged, then split between positive staining and isotype controls. Extracellular antibodies were incubated with cells for 30 min: CD45-APC/Cy7 (clone 30-F11, BioLegend, 1:200) for leukocytes, CD11b-PerCP-Cy5.5 (clone M1/70, BioLegend, 1:200) for myeloid cells, Ly6G-Pacific Blue (clone 1A8, BioLegend, 1:200) or its isotype (Rat IgG2a, BioLegend, 1:200) for neutrophils, and Goat anti-Human/Mouse myeloperoxidase/MPO (AF3667, R&D systems, 1:50) or no antibody for myeloperoxidase. Cells were diluted with NDS-FACS buffer and centrifuged twice. For the MPO antibody, all cells were incubated with donkey anti-goat Alexa Fluor 647 (A-21447, Invitrogen, 1:2000) for 30 min, then washed with NDS-FACS buffer and centrifuged twice. Cells were fixed with Cyto-Fast Fix Perm solution (BioLegend) for 20 min, then washed according to manufacturer’s instructions. For overnight storage at 4°C, cells were centrifuged and resuspended in NDS-FACS buffer. The next day, cells were resuspended in Cyto-Fast Perm Wash buffer (BioLegend) and centrifuged. Intracellular staining for citrullinated histones was accomplished with rabbit anti-CitH3 antibody (Abcam, 1:2000 in 100 µL Perm Wash buffer) or a no antibody control incubation for 30 min. Cells were resuspended in NDS-FACS buffer and centrifuged twice. CitH3 was visualized with donkey anti-rabbit Alexa Fluor 488 (Invitrogen, 1:2000 in NDS-FACS buffer), 30 min incubation. Cells were washed with NDS-FACS buffer and resuspended prior to analysis. Cells were assessed with a BD Fortessa X-20 flow cytometer and data were analyzed with FlowJo software.

### ETs Capture ELISA

ET complexes were quantified as previously described^28^. Briefly, a Nunc uncoated 96-well plate was coated with CitH3 capture antibody (Abcam, 1:250) diluted in coating buffer (15mM Na_2_CO_3_, 35mM NaHCO_3_ in 1X PBS) and incubated overnight at 4°C. Plates were washed with 1X PBS (x3), then blocked with 5% Bovine Serum Albumin (BSA) for 2 hours at RT. Cell-free supernatant of spinal cord and blood were diluted with 1% BSA in PBS and incubated on plate overnight at 4°C. Following this, the plate was washed (1% BSA, 0.05% Tween-20 in 1X PBS, 3×200 µL) three times, then incubated for 2 hrs at RT with anti-DNA-POD detection antibody (Roche, 1:100) according to manufacturer instructions. Plate was washed with wash buffer three times, then incubated with room temperature TMB substrate solution (1X, Invitrogen) in the dark at RT for about 10 min. The reaction was quenched with 0.5 M phosphoric acid. The absorbance of each well was taken at 450 nm.

### White Matter Sparing

White matter sparing was assessed via Eriochrome cyanine staining as previously described^28^. Tissue sections were cleared in Citrisolv (5 min, Decon Labs), then hydrated in a series of ethanol baths (100% x 2, 95%, 70%, 50%, 1 min) followed by distilled water (2 min). Myelin was stained with Eriochrome Cyanine R solution (0.2% Eriochrome Cyanine R, 90 mM H_2_SO_4_, 5.6% FeCl_3_ in dH_2_O) for 10 min followed by three water washes (ddH_2_O, 1 min). Sections were differentiated with 5.6% FeCl_3_ in water for 3 min, then washed in ddH_2_O three times. Sections were rehydrated in a series of ethanol baths (70%, 1 min; 95%, 2 min; 100%, 1 min x 2), then cleared with Citrisolv (5 min x 2). Slides were airdried, then coverslipped with Cytoseal. Images were acquired with a 10 X objective on the Leica DM6B upright microscope (RRID: SCR_022128). Tissue sparing was analyzed as previously described using ImageJ—total section area and white matter area were quantified in tissue sections 250 µm apart spanning the lesion site. The section with the lowest percentage of white matter was identified as the lesion center.

### Single Cell RNA Sequencing Analysis

Single-cell RNA sequencing (scRNA-seq) datasets (accessions SRR17591922–SRR17591925) were downloaded from the Sequence Read Archive (SRA) using the NCBI SRA Toolkit (version 3.0). Raw files were converted to FASTQ format, sequencing data were aligned to the mouse transcriptome reference (GRCm39-2024-A), and unique molecular identifiers (UMIs) were quantified using Cell Ranger software (version 9.0.1). The resulting feature-barcode matrices were analyzed using the Seurat package (version 4.3) in R (version 4.2.2). Quality control filters removed cells expressing fewer than 200 or more than 4000 genes, as well as cells with mitochondrial genes above 2%. Cells were clustered using the shared nearest neighbor algorithm and visualized via Uniform Manifold Approximation and Projection (UMAP). Sex classification of neutrophils was based on expression of the female-specific marker Xist and the male-specific marker Ddx3y^38^. Differential expression analysis comparing male and female neutrophils was performed for the Padi gene family (Padi1, Padi2, Padi3, Padi4, and Padi6).

### Statistics

Statistics were analyzed using GraphPad Prism 10. Flow cytometry, ELISA, tissue sparing, and LRWT data were analyzed as follows: for pairwise comparisons, an F test to compare variances was performed, followed by an unpaired two-tailed t-test. If variances were significantly different, Welsh’s correction was used. For comparisons between groups with two or more factors (i.e., genotype and sex), a two-way ANOVA followed by Tukey’s post-hoc test for multiple comparisons was performed. BMS and tissue sparing were analyzed as follows: for comparisons between groups with three or more factors (i.e., treatment, sex, and time), a three-way repeated measures ANOVA or appropriate mixed-effects model was used followed by Tukey’s post-hoc test for multiple comparisons. For scRNAseq, transcript level comparisons were made using the Mann-Whitney U test.

#### Exclusions

For data from ELISA and FC experiments, statistical outliers were removed using Grubbs’ test (alpha=0.05). For short term studies, animals with non-standard surgeries (abnormal impact force or displacement) were excluded from analyses. For long-term studies, data from animals that had an abnormal impact force or displacement, died during the study, or exhibited a BMS score > 2 at 1 dpi were excluded from analyses. Excluded data is retained in the data repository associated with this manuscript.

## Funding

This work was supported by funding from The Craig H. Neilsen Foundation Grant #648714, Mission Connect (a program of TIRR Foundation), and NIH R01NS122961.

## Declaration of Interest

The authors declare that they have no known competing financial interests or personal relationships that could have appeared to influence the work reported in this paper.

## Acknowledgements

The authors would like to thank Texas A&M University Microscopy and Imaging Center (RRID: SCR_022128) and Texas A&M University Flow Cytometry & Cell Sorting Facility for the use of their equipment. The diagrams in Figures 1A and S1A were created using Biorender.com.

## CRediT Authorship Statement

**Shelby Reid:** Conceptualization, Methodology, Validation, Formal analysis, Investigation, Visualization, Data Curation, Writing - Original Draft, Project administration. **Ashley Tran:** Validation, Investigation. **Miranda Leal-Garcia**: Methodology, Investigation. **Sachit Devaraj:** Investigation. **Mustafa Ozturgut:** Data Curation, Formal Analysis, Visualization. **Dylan McCreedy:** Writing – review & editing, Resources, Supervision, Project administration, Funding acquisition, Conceptualization

## Data availability

All associated data from this manuscript will be deposited in the Open Data Commons for Spinal Cord Injury (ODC-SCI) upon publication (https://odc-sci.org/).

## Supplemental Figures

**Figure S1:**
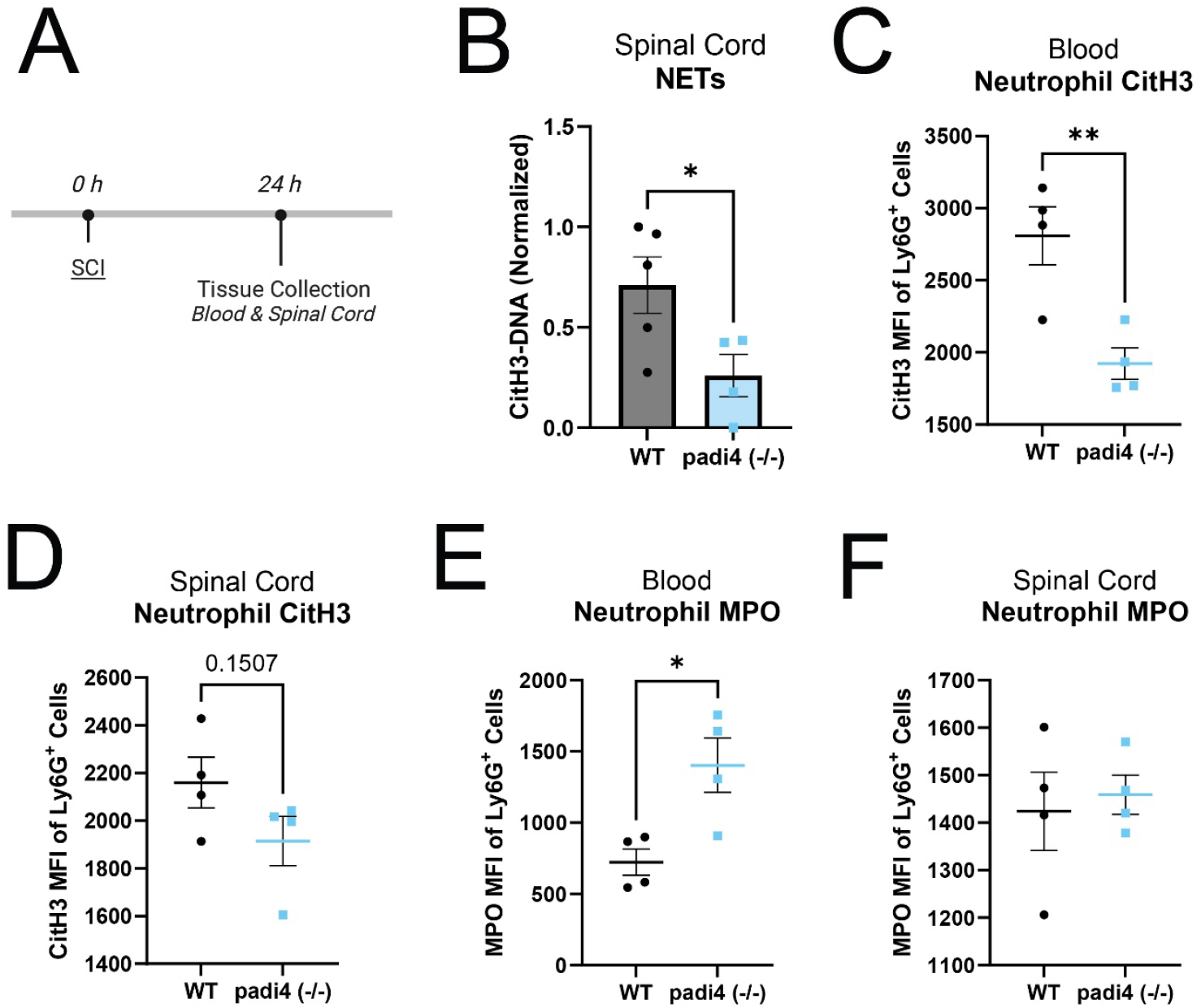
PAD4 KO alters neutrophil effector function at 24 hours after SCI. **(A)** At 24 hr following moderate T9 contusion SCI, *Padi4* (-/-) and WT mouse spinal cords and blood were collected and assessed for ETosis via flow cytometry and capture ELISA. **(B)** Extracellular trap (ET) complexes at the spinal cord injury site were reduced in *Padi4(-/-)* mice compared to littermate controls (Unpaired Student’s *t*-test, n=4-5, p=0.0440). **(C-D)** Intracellular citrullination of histone 3 (CitH3) in **(C)** blood neutrophils was reduced after injury compared to littermate controls (Unpaired student’s *t*-test, n=4, p=0.0083). A similar trend was observed in **(D)** spinal cord neutrophils (Unpaired student’s *t*-test, n=4, p=0.1507). Data shown as mean ± SEM. *p<0.05, **p<0.01. **(E-F)** Extracellular myeloperoxidase (MPO) was elevated on **(E)** blood neutrophils while no change was observed in **(F)** spinal cord neutrophils. (Unpaired t-test; n=4, p_blood_=0.0185, p_spinalcord_= 0.7170).

## Notes

### Competing Interest Statement

The authors have declared no competing interest.

## References

1. Hellenbrand DJ, Quinn CM, Piper ZJ, et al. Inflammation after spinal cord injury: a review of the critical timeline of signaling cues and cellular infiltration. J Neuroinflammation 2021;18(1):284; doi: 10.1186/s12974-021-02337-2.

2. Beck KD, Nguyen HX, Galvan MD, et al. Quantitative analysis of cellular inflammation after traumatic spinal cord injury: evidence for a multiphasic inflammatory response in the acute to chronic environment. Brain 2010;133(Pt 2):433–447; doi: 10.1093/brain/awp322.

3. Jin L-Y, Li J, Wang K-F, et al. Blood–Spinal Cord Barrier in Spinal Cord Injury: A Review. Journal of Neurotrauma 2021;38(9):1203–1224; doi: 10.1089/neu.2020.7413.

4. Stirling DP, Yong VW. Dynamics of the inflammatory response after murine spinal cord injury revealed by flow cytometry. J Neurosci Res 2008;86(9):1944–1958; doi: 10.1002/jnr.21659.

5. Zivkovic S, Ayazi M, Hammel G, et al. For Better or for Worse: A Look Into Neutrophils in Traumatic Spinal Cord Injury. Front Cell Neurosci 2021;15:648076; doi: 10.3389/fncel.2021.648076.

6. Neirinckx V, Coste C, Franzen R, et al. Neutrophil contribution to spinal cord injury and repair. J Neuroinflammation 2014;11(1):150; doi: 10.1186/s12974-014-0150-2.

7. Brennan FH, Jogia T, Gillespie ER, et al. Complement receptor C3aR1 controls neutrophil mobilization following spinal cord injury through physiological antagonism of CXCR2. JCI Insight 2019;4(9); doi: 10.1172/jci.insight.98254.

8. Jogia T, Lübstorf T, Jacobson E, et al. Prognostic value of early leukocyte fluctuations for recovery from traumatic spinal cord injury. Clin Transl Med 2021;11(1):e272; doi: 10.1002/ctm2.272.

9. Saiwai H, Ohkawa Y, Yamada H, et al. The LTB4-BLT1 axis mediates neutrophil infiltration and secondary injury in experimental spinal cord injury. Am J Pathol 2010;176(5):2352–2366; doi: 10.2353/ajpath.2010.090839.

10. Poli V, Zanoni I. Neutrophil intrinsic and extrinsic regulation of NETosis in health and disease. Trends in Microbiology 2023;31(3):280–293; doi: 10.1016/j.tim.2022.10.002.

11. Thiam HR, Wong SL, Wagner DD, et al. Cellular Mechanisms of NETosis. Annu Rev Cell Dev Biol 2020; doi: 10.1146/annurev-cellbio-020520-111016.

12. Ma Q, Steiger S. Neutrophils and extracellular traps in crystal-associated diseases. Trends Mol Med 2024;30(9):809–823; doi: 10.1016/j.molmed.2024.05.010.

13. Etulain J, Martinod K, Wong SL, et al. P-selectin promotes neutrophil extracellular trap formation in mice. Blood 2015;126(2):242–246; doi: 10.1182/blood-2015-01-624023.

14. Vaibhav K, Braun M, Alverson K, et al. Neutrophil extracellular traps exacerbate neurological deficits after traumatic brain injury. Sci Adv 2020;6(22):eaax8847; doi: 10.1126/sciadv.aax8847.

15. Krämer TJ, Pickart F, Pöttker B, et al. Early DNase-I therapy delays secondary brain damage after traumatic brain injury in adult mice. Sci Rep 2023;13(1):4348; doi: 10.1038/s41598-023-30421-5.

16. Liu Y-W, Zhang J, Bi W, et al. Histones of Neutrophil Extracellular Traps Induce CD11b Expression in Brain Pericytes Via Dectin-1 after Traumatic Brain Injury. Neurosci Bull 2022;38(10):1199–1214; doi: 10.1007/s12264-022-00902-0.

17. Cao Y, Shi M, Liu L, et al. Inhibition of neutrophil extracellular trap formation attenuates NLRP1-dependent neuronal pyroptosis via STING/IRE1α pathway after traumatic brain injury in mice. Frontiers in Immunology 2023;14.

18. Shi G, Liu L, Cao Y, et al. Inhibition of neutrophil extracellular trap formation ameliorates neuroinflammation and neuronal apoptosis via STING-dependent IRE1α/ASK1/JNK signaling pathway in mice with traumatic brain injury. Journal of Neuroinflammation 2023;20(1):222; doi: 10.1186/s12974-023-02903-w.

19. Mi L, Min X, Shi M, et al. Neutrophil extracellular traps aggravate neuronal endoplasmic reticulum stress and apoptosis via TLR9 after traumatic brain injury. Cell Death Dis 2023;14(6):1–12; doi: 10.1038/s41419-023-05898-7.

20. Mu Q, Yao K, Syeda MZ, et al. Neutrophil Targeting Platform Reduces Neutrophil Extracellular Traps for Improved Traumatic Brain Injury and Stroke Theranostics. Advanced Science 2024;11(21):2308719; doi: 10.1002/advs.202308719.

21. Li B, Xu L, Wang Z, et al. Neutrophil Extracellular Traps Regulate Surgical Brain Injury by Activating the cGAS-STING Pathway. Cell Mol Neurobiol 2024;44(1):36; doi: 10.1007/s10571-024-01470-9.

22. Jin Z, Sun J, Song Z, et al. Neutrophil extracellular traps promote scar formation in post-epidural fibrosis. npj Regen Med 2020;5(1):1–9; doi: 10.1038/s41536-020-00103-1.

23. Hua F, Wang H-R, Bai Y-F, et al. Substance P promotes epidural fibrosis via induction of type 2 macrophages. Neural Regeneration Research 2023;18(10):2252; doi: 10.4103/1673-5374.369120.

24. Feng Z, Min L, Liang L, et al. Neutrophil Extracellular Traps Exacerbate Secondary Injury via Promoting Neuroinflammation and Blood–Spinal Cord Barrier Disruption in Spinal Cord Injury. Frontiers in Immunology 2021;12.

25. Zhang C, Guo D, Qiao H, et al. Macrophage Extracellular Traps Exacerbate Secondary Spinal Cord Injury by Modulating Macrophage/Microglia Polarization via LL37/P2×7R/NF-κB Signaling Pathway. Oxidative Medicine and Cellular Longevity 2022;2022:e9197940; doi: 10.1155/2022/9197940.

26. Morishima Y, Kawabori M, Yamazaki K, et al. Intravenous Administration of Mesenchymal Stem Cell-Derived Exosome Alleviates Spinal Cord Injury by Regulating Neutrophil Extracellular Trap Formation through Exosomal miR-125a-3p. Int J Mol Sci 2024;25(4):2406; doi: 10.3390/ijms25042406.

27. Tang C, Jin Y, Wu M, et al. A biomimic anti-neuroinflammatory nanoplatform for active neutrophil extracellular traps targeting and spinal cord injury therapy. Mater Today Bio 2024;28:101218; doi: 10.1016/j.mtbio.2024.101218.

28. Reid SK, Leal-Garcia ME, Tran AV, et al. Recombinant human DNase treatment mitigates extracellular trap mediated damage and improves long-term recovery after spinal cord injury in male mice. Brain Behav Immun 2025;128:456–468; doi: 10.1016/j.bbi.2025.04.033.

29. Zhou Y, Chen B, Mittereder N, et al. Spontaneous Secretion of the Citrullination Enzyme PAD2 and Cell Surface Exposure of PAD4 by Neutrophils. Front Immunol 2017;8:1200; doi: 10.3389/fimmu.2017.01200.

30. Wu Z, Deng Q, Pan B, et al. Inhibition of PAD2 Improves Survival in a Mouse Model of Lethal LPS-Induced Endotoxic Shock. Inflammation 2020;43(4):1436–1445; doi: 10.1007/s10753-020-01221-0.

31. Domiciano TP, Lee Y, Carvalho TT, et al. Redundant role of PAD2 and PAD4 in the development of cardiovascular lesions in a mouse model of Kawasaki disease vasculitis. Clinical and Experimental Immunology 2024;218(3):314–328; doi: 10.1093/cei/uxae080.

32. Yu X, Song Y, Dong T, et al. Loss of PADI2 and PADI4 ameliorates sepsis-induced acute lung injury by suppressing NLRP3+ macrophages. JCI Insight 2024;9(22):e181686; doi: 10.1172/jci.insight.181686.

33. Luo Y, Arita K, Bhatia M, et al. Inhibitors and inactivators of protein arginine deiminase 4: functional and structural characterization. Biochemistry 2006;45(39):11727–11736; doi: 10.1021/bi061180d.

34. Mansouri P, Mansouri P, Behmard E, et al. Peptidylarginine deiminase (PAD): A promising target for chronic diseases treatment. Int J Biol Macromol 2024;278(Pt 3):134576; doi: 10.1016/j.ijbiomac.2024.134576.

35. Jo J, Hugonnet H, Lee MJ, et al. Digital Cytometry: Extraction of Forward and Side Scattering Signals From Holotomography. Journal of Biophotonics 2025;n/a(n/a):e202400387; doi: 10.1002/jbio.202400387.

36. Wu T, Tan JHL, Sin W, et al. Cell Granularity Reflects Immune Cell Function and Enables Selection of Lymphocytes with Superior Attributes for Immunotherapy. Adv Sci (Weinh) 2023;10(28):2302175; doi: 10.1002/advs.202302175.

37. Fernández-Segura E, García JM, Santos JL, et al. Shape, F-actin, and surface morphology changes during chemotactic peptide-induced polarity in human neutrophils. Anat Rec 1995;241(4):519–528; doi: 10.1002/ar.1092410410.

38. Kim M, Lu RJ, Benayoun BA. Single-cell RNA-seq of primary bone marrow neutrophils from female and male adult mice. Sci Data 2022;9(1):442; doi: 10.1038/s41597-022-01544-7.

39. Michel-Flutot P, Bourcier CH, Emam L, et al. Extracellular traps formation following cervical spinal cord injury. European Journal of Neuroscience 2023;57(4):692–704; doi: 10.1111/ejn.15902.

40. McCreedy DA, Abram CL, Hu Y, et al. Spleen tyrosine kinase facilitates neutrophil activation and worsens long-term neurologic deficits after spinal cord injury. Journal of Neuroinflammation 2021;18(1):302; doi: 10.1186/s12974-021-02353-2.

41. Nauseef WM, Kubes P. Pondering neutrophil extracellular traps with healthy skepticism. Cell Microbiol 2016;18(10):1349–1357; doi: 10.1111/cmi.12652.

42. Branzk N, Lubojemska A, Hardison SE, et al. Neutrophils sense microbe size and selectively release neutrophil extracellular traps in response to large pathogens. Nat Immunol 2014;15(11):1017–1025; doi: 10.1038/ni.2987.

43. Hampson P, Dinsdale RJ, Wearn CM, et al. Neutrophil Dysfunction, Immature Granulocytes, and Cell-free DNA are Early Biomarkers of Sepsis in Burn-injured Patients: A Prospective Observational Cohort Study. Ann Surg 2017;265(6):1241–1249; doi: 10.1097/SLA.0000000000001807.

44. Zhang D, Chen G, Manwani D, et al. Neutrophil ageing is regulated by the microbiome. Nature 2015;525(7570):528–532; doi: 10.1038/nature15367.

45. Ortmann W, Kolaczkowska E. Age is the work of art? Impact of neutrophil and organism age on neutrophil extracellular trap formation. Cell Tissue Res 2018;371(3):473–488; doi: 10.1007/s00441-017-2751-4.

46. Barth CR, Luft C, Funchal GA, et al. LPS-induced neonatal stress in mice affects the response profile to an inflammatory stimulus in an age and sex-dependent manner. Developmental Psychobiology 2016;58(5):600–613; doi: 10.1002/dev.21404.

47. Lu RJ, Taylor S, Contrepois K, et al. Multi-omic profiling of primary mouse neutrophils predicts a pattern of sex- and age-related functional regulation. Nat Aging 2021;1(8):715–733; doi: 10.1038/s43587-021-00086-8.

48. Hazeldine J, Harris P, Chapple IL, et al. Impaired neutrophil extracellular trap formation: a novel defect in the innate immune system of aged individuals. Aging Cell 2014;13(4):690–698; doi: 10.1111/acel.12222.

49. Ishikawa M, Murakami H, Higashi H, et al. Sex Differences of Neutrophil Extracellular Traps on Lipopolysaccharide-Stimulated Human Neutrophils. Surgical Infections 2024;25(7):505–512; doi: 10.1089/sur.2024.016.

50. Patel D, Dodd WS, Lucke-Wold B, et al. Neutrophils: Novel Contributors to Estrogen-Dependent Intracranial Aneurysm Rupture Via Neutrophil Extracellular Traps. J Am Heart Assoc 2023;12(21):e029917; doi: 10.1161/JAHA.123.029917.

51. Jarrot P-A, Kim J, Chan W, et al. Sex-specific NLRP3 activation in neutrophils promotes neutrophil recruitment and NETosis in the murine model of diffuse alveolar hemorrhage. Front Immunol 2024;15:1466234; doi: 10.3389/fimmu.2024.1466234.

52. Wang L, Huang F-Y, Dai S-Z, et al. Progesterone modulates the immune microenvironment to suppress ovalbumin-induced airway inflammation by inhibiting NETosis. Sci Rep 2024;14(1):17241; doi: 10.1038/s41598-024-66439-6.

53. Melo Z, Gutierrez-Mercado YK, Garcia-Martínez D, et al. Sex-dependent mechanisms involved in renal tolerance to ischemia-reperfusion: Role of inflammation and histone H3 citrullination. Transpl Immunol 2020;63:101331; doi: 10.1016/j.trim.2020.101331.

54. Giaglis S, Stoikou M, Sur Chowdhury C, et al. Multimodal Regulation of NET Formation in Pregnancy: Progesterone Antagonizes the Pro-NETotic Effect of Estrogen and G-CSF. Front Immunol 2016;7:565; doi: 10.3389/fimmu.2016.00565.

55. Dai S-Z, Wu R-H, Chen H, et al. Progesterone suppresses rhinovirus-induced airway inflammation by inhibiting neutrophil infiltration and extracellular traps formation. Int Immunopharmacol 2025;144:113714; doi: 10.1016/j.intimp.2024.113714.

56. Poole JA, Thiele GM, Ramler E, et al. Combined repetitive inhalant endotoxin and collagen-induced arthritis drive inflammatory lung disease and arthritis severity in a testosterone-dependent manner. Am J Physiol Lung Cell Mol Physiol 2024;326(3):L239–L251; doi: 10.1152/ajplung.00221.2023.

57. Yasuda H, Sonoda A, Yamamoto M, et al. 17-β-estradiol enhances neutrophil extracellular trap formation by interaction with estrogen membrane receptor. Arch Biochem Biophys 2019;663:64–70; doi: 10.1016/j.abb.2018.12.028.

58. Mohanan S, Horibata S, McElwee JL, et al. Identification of macrophage extracellular trap-like structures in mammary gland adipose tissue: a preliminary study. Front Immunol 2013;4:67; doi: 10.3389/fimmu.2013.00067.

59. Ghasempour G, Zamani-Garmsiri F, Shojaei S, et al. Vitamin D3 and estradiol alter PAD2 expression and activity levels in C6 glioma cells. Mult Scler Relat Disord 2021;56:103221; doi: 10.1016/j.msard.2021.103221.

60. Dong S, Zhang Z, Takahara H. Estrogen-enhanced peptidylarginine deiminase type IV gene (PADI4) expression in MCF-7 cells is mediated by estrogen receptor-alpha-promoted transfactors activator protein-1, nuclear factor-Y, and Sp1. Mol Endocrinol 2007;21(7):1617–1629; doi: 10.1210/me.2006-0550.

61. Nagata S, Yamagiwa M, Inoue K, et al. Estrogen regulates peptidylarginine deiminase levels in a rat pituitary cell line in culture. J Cell Physiol 1990;145(2):333–339; doi: 10.1002/jcp.1041450219.

62. Horibata S, Coonrod SA, Cherrington BD. Role for peptidylarginine deiminase enzymes in disease and female reproduction. J Reprod Dev 2012;58(3):274–282; doi: 10.1262/jrd.2011-040.

63. Christensen AO, Li G, Young CH, et al. Peptidylarginine deiminase enzymes and citrullinated proteins in female reproductive physiology and associated diseases†. Biology of Reproduction 2022;107(6):1395–1410; doi: 10.1093/biolre/ioac173.

64. Stewart AN, MacLean SM, Stromberg AJ, et al. Considerations for Studying Sex as a Biological Variable in Spinal Cord Injury. Front Neurol 2020;11:802; doi: 10.3389/fneur.2020.00802.

65. Späni CB, Braun DJ, Van Eldik LJ. Sex-related responses after traumatic brain injury: Considerations for preclinical modeling. Frontiers in Neuroendocrinology 2018;50:52–66; doi: 10.1016/j.yfrne.2018.03.006.

66. Gupte R, Brooks W, Vukas R, et al. Sex Differences in Traumatic Brain Injury: What We Know and What We Should Know. J Neurotrauma 2019;36(22):3063–3091; doi: 10.1089/neu.2018.6171.

67. Kim T, Chelluboina B, Chokkalla AK, et al. Age and sex differences in the pathophysiology of acute CNS injury. Neurochem Int 2019;127:22–28; doi: 10.1016/j.neuint.2019.01.012.

68. Stewart AN, Lowe JL, Glaser EP, et al. Acute inflammatory profiles differ with sex and age after spinal cord injury. J Neuroinflammation 2021;18(1):113; doi: 10.1186/s12974-021-02161-8.

69. Stewart AN, Glaser EP, Bailey WM, et al. Immunoglobulin G Is Increased in the Injured Spinal Cord in a Sex- and Age-Dependent Manner. J Neurotrauma 2022;39(15–16):1090–1098; doi: 10.1089/neu.2022.0011.

70. Ghosh M, Lee J, Burke AN, et al. Sex Dependent Disparities in the Central Innate Immune Response after Moderate Spinal Cord Contusion in Rat. Cells 2024;13(7):645; doi: 10.3390/cells13070645.

71. Li Y, Ritzel RM, Lei Z, et al. Sexual dimorphism in neurological function after SCI is associated with disrupted neuroinflammation in both injured spinal cord and brain. Brain Behav Immun 2022;101:1–22; doi: 10.1016/j.bbi.2021.12.017.

72. Falcão AM, Meijer M, Scaglione A, et al. PAD2-Mediated Citrullination Contributes to Efficient Oligodendrocyte Differentiation and Myelination. Cell Reports 2019;27(4):1090-1102.e10; doi: 10.1016/j.celrep.2019.03.108.

73. Boeltz S, Amini P, Anders H-J, et al. To NET or not to NET:current opinions and state of the science regarding the formation of neutrophil extracellular traps. Cell Death Differ 2019;26(3):395–408; doi: 10.1038/s41418-018-0261-x.

74. Jorch SK, Kubes P. An emerging role for neutrophil extracellular traps in noninfectious disease. Nature Medicine 2017;23(3):279–287; doi: 10.1038/nm.4294.

75. Castanheira FVS, Kubes P. Neutrophils and NETs in modulating acute and chronic inflammation. Blood 2019;133(20):2178–2185; doi: 10.1182/blood-2018-11-844530.

76. Warnatsch A, Ioannou M, Wang Q, et al. Inflammation. Neutrophil extracellular traps license macrophages for cytokine production in atherosclerosis. Science 2015;349(6245):316–320; doi: 10.1126/science.aaa8064.

77. Kenny EF, Herzig A, Krüger R, et al. Diverse stimuli engage different neutrophil extracellular trap pathways. Brakhage AA. ed. eLife 2017;6:e24437; doi: 10.7554/eLife.24437.

78. Guiducci E, Lemberg C, Küng N, et al. Candida albicans-Induced NETosis Is Independent of Peptidylarginine Deiminase 4. Front Immunol 2018;9; doi: 10.3389/fimmu.2018.01573.

79. Byun DJ, Lee J, Ko K, et al. NLRP3 exacerbates EAE severity through ROS-dependent NET formation in the mouse brain. Cell Commun Signal 2024;22(1):1–16; doi: 10.1186/s12964-023-01447-z.

80. Sebina I, Rashid RB, Sikder MAA, et al. IFN-λ Diminishes the Severity of Viral Bronchiolitis in Neonatal Mice by Limiting NADPH Oxidase-Induced PAD4-Independent NETosis. J Immunol 2022;208(12):2806–2816; doi: 10.4049/jimmunol.2100876.

81. Díaz-Godínez C, Fonseca Z, Néquiz M, et al. Entamoeba histolytica Trophozoites Induce a Rapid Non-classical NETosis Mechanism Independent of NOX2-Derived Reactive Oxygen Species and PAD4 Activity. Front Cell Infect Microbiol 2018;8:184; doi: 10.3389/fcimb.2018.00184.

82. Wu S-Y, Weng C-L, Jheng M-J, et al. Candida albicans triggers NADPH oxidase-independent neutrophil extracellular traps through dectin-2. PLoS Pathog 2019;15(11):e1008096; doi: 10.1371/journal.ppat.1008096.

83. Spengler J, Lugonja B, Jimmy Ytterberg A, et al. Release of Active Peptidyl Arginine Deiminases by Neutrophils Can Explain Production of Extracellular Citrullinated Autoantigens in Rheumatoid Arthritis Synovial Fluid. Arthritis & Rheumatology 2015;67(12):3135–3145; doi: 10.1002/art.39313.

84. Zheng L, Nagar M, Maurais AJ, et al. Calcium Regulates the Nuclear Localization of Protein Arginine Deiminase 2. Biochemistry 2019;58(27):3042–3056; doi: 10.1021/acs.biochem.9b00225.

85. Holmes CL, Shim D, Kernien J, et al. Insight into Neutrophil Extracellular Traps through Systematic Evaluation of Citrullination and Peptidylarginine Deiminases. J Immunol Res 2019;2019:2160192; doi: 10.1155/2019/2160192.

86. Bawadekar M, Shim D, Johnson CJ, et al. Peptidylarginine deiminase 2 is required for tumor necrosis factor alpha-induced citrullination and arthritis, but not neutrophil extracellular trap formation. Journal of Autoimmunity 2017;80:39–47; doi: 10.1016/j.jaut.2017.01.006.

87. Stachowicz A, Pandey R, Sundararaman N, et al. Protein arginine deiminase 2 (PAD2) modulates the polarization of THP-1 macrophages to the anti-inflammatory M2 phenotype. J Inflamm (Lond) 2022;19(1):20; doi: 10.1186/s12950-022-00317-8.

88. Tian Y, Qu S, Alam HB, et al. Peptidylarginine deiminase 2 has potential as both a biomarker and therapeutic target of sepsis. JCI Insight 2020;5(20):e138873. 138873; doi: 10.1172/jci.insight.138873.

89. Bashar SJ, Holmes CL, Shelef MA. Macrophage extracellular traps require peptidylarginine deiminase 2 and 4 and are a source of citrullinated antigens bound by rheumatoid arthritis autoantibodies. Front Immunol 2024;15:1167362; doi: 10.3389/fimmu.2024.1167362.

90. Chen T, Wang Y, Nan Z, et al. Interaction Between Macrophage Extracellular Traps and Colon Cancer Cells Promotes Colon Cancer Invasion and Correlates With Unfavorable Prognosis. Front Immunol 2021;12:779325; doi: 10.3389/fimmu.2021.779325.

91. Anonymous. 030315 - Pad4[-] Strain Details. n.d. Available from: https://www.jax.org/strain/030315 [Last accessed: 12/23/2024].

92. Cummings BJ, Engesser-Cesar C, Cadena G, et al. Adaptation of a ladder beam walking task to assess locomotor recovery in mice following spinal cord injury. Behav Brain Res 2007;177(2):232–241; doi: 10.1016/j.bbr.2006.11.042.

